# Development and validation of a reliable DNA copy-number-based machine learning algorithm (*CopyClust*) for breast cancer integrative cluster classification

**DOI:** 10.1101/2023.11.21.568129

**Authors:** Cameron C. Young, Katherine Eason, Raquel Manzano Garcia, Richard Moulange, Sach Mukherjee, Suet-Feung Chin, Carlos Caldas, Oscar M. Rueda

## Abstract

The Integrative Clusters (IntClusts) provide a framework for the classification of breast cancer tumors into 10 distinct genomic subtypes based on DNA copy number and gene expression. Current classifiers achieve only low accuracy without gene expression data, warranting the development of new approaches to copy-number-only-based IntClust classification. A novel XGBoost-driven classification algorithm, *CopyClust*, was trained using genomic features from METABRIC and validated on TCGA achieving a nine-percentage point or greater improvement in overall IntClust subtype classification accuracy.

## Introduction

Extensive copy number (CN) and gene expression (GE) integrated analysis of the METABRIC cohort revealed 10 unique breast cancer tumor subtypes (Integrative Clusters [IntClusts]) each with characteristic genomic and transcriptomic architecture and genomic driver events^1,2^. Clinically, the IntClusts showed distinct patterns of survival^1^ and risk of disease relapse^3^. Crucially, this work reveals that by integrating multi-omic features, it is possible to derive more robust patient classifiers that can stratify tumours based on candidate genomic driver events, as opposed to GE-only approaches, like PAM50^4^. Furthermore, it provides a framework for personalized breast cancer therapeutic strategies with extensive clinical utility^5,6^. For clinical application, new samples, which are initially unlabelled, must be assigned to a class. However, current approaches to classify unlabelled tumor samples into IntClusts rely on using a combined CN and GE-focused approach^7^, limiting classification accuracy among cancer samples without transcriptomic data. With the increase in cancer genomics consortia and widespread availability of public data without GE profiling, a novel method to subtype tumors from independent cohorts based on CN data alone is warranted. Here, we present the development and validation of a reliable, flexible, platform-independent CN-driven machine learning algorithm (*CopyClust*) for IntClust classification as an open-source R package.

## Methods

The 1,980 breast cancer samples from METABRIC (internal validation) and the 1,075 samples from TCGA (external validation) with available CN and GE data were used to train and validate multiclass hyperparameter-optimized XGBoost^8^ machine learning algorithms. For METABRIC, IntClust label was assigned from the original manuscript^1^, while for TCGA, label was assigned via the *iC10* classifier^7^ using CN and GE data. To reduce noise, a piecewise constant fit (PCF) was applied to the CN profiles of the METABRIC samples in each IntClust and unique breakpoints across the 10 profiles were identified and converted into 478 genomic regions. The mean CN in each region was calculated, and these were used as features for XGBoost models. METABRIC samples were split into a training cohort (80%) and validation cohort (20%). Intra-IntClust outliers were identified via local outlier factor (LOF) and removed from the training cohort prior to model training.

Two model approaches were implemented: a 10-class model with the final IntClust label predicted by a single multiclass model and a 6-class model with binary reclassification in which four pairs of IntClusts were combined for initial multiclass classification, then assigned an IntClust label based on the prediction of a second binary classifier trained on that pair. These pairs were selected based on their similar CN profiles and frequent misclassification in the 10-class model. The 6-class model with binary reclassification model was selected for final model implementation due to superior performance (**Supplementary Table S1**). Scaling of feature values was performed prior to model implementation on external cohorts. Model performance was internally validated on 392 (20%) held-out METABRIC samples and externally validated on the TCGA dataset, with CN data generated from both single nucleotide polymorphism (SNP) arrays and whole exome sequencing (WES) (**Figure 1**). More details of the methodology used and results for this study can be found in the **Supplemental Methods** and **Supplemental Tables S1–S7 and Supplemental Figures S1–S18**.

**Figure 1.**
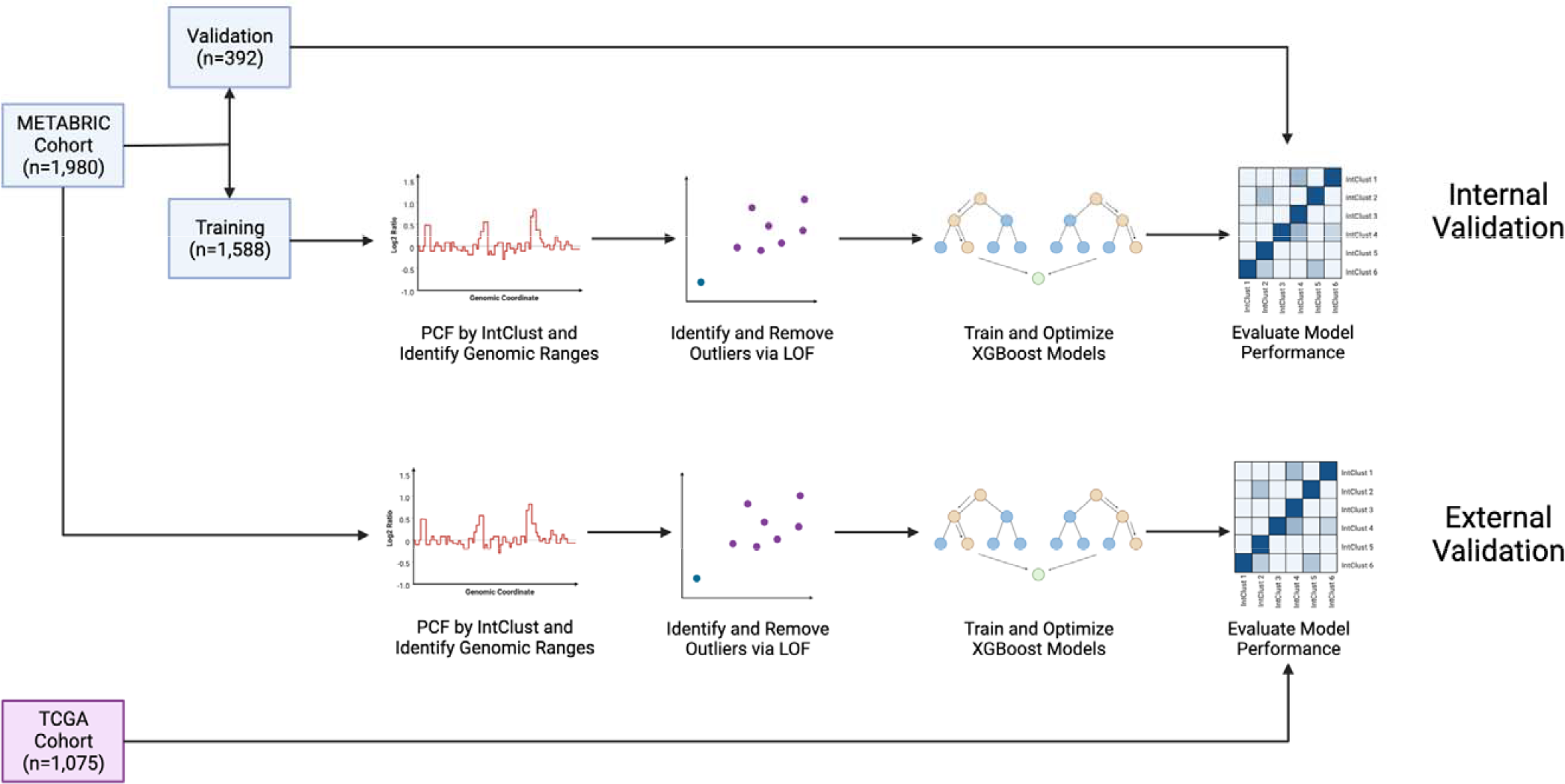
Workflow of algorithm development for internal and external validation.

## Results and Discussion

When compared to other hyperparameter-optimized classifier algorithms including Random Forest, Support Vector Machine, LightGBM, and Prediction Analysis of Microarrays, XGBoost performed best in terms of overall recall and Matthews Correlation Coefficient (MCC) (**Table 1**) and was selected as the approach for *CopyClust. CopyClust* achieved high classification performance across both the TCGA SNP and WES datasets (**Table 1** and **Figure 2**). The classifier produced an overall recall of 81%, precision of 82%, and balanced accuracy of 89% when applied to the TCGA SNP dataset with an MCC of 0.787. Applied to the TCGA WES dataset, *CopyClust* produced an overall recall of 79%, precision of 80%, balanced accuracy of 88%, and MCC of 0.759. Across both datasets, IntClust 3 and IntClust 8 were the most misclassified pair of IntClusts, likely due to their similar CN profiles. IntClust 6 experienced the lowest individual recall, which could be due to differences in the distribution of IntClusts between cohorts (**Supplementary Figure S2**), leading to model miscalibration^9^.

**Table 1.** Overall recall of different classifier approaches

| Model Approach<sup>a</sup> | Overall Recall (95% Confidence Interval)<sup>b</sup> | Matthews Correlation Coefficient |
| --- | --- | --- |
| Random Forest | 68.4% (65.5%, 71.1%) | 0.651 |
| Support Vector Machine | 68.6% (65.7%, 71.3%) | 0.645 |
| LightGBM | 71.4% (68.5%, 74.0%) | 0.677 |
| Prediction Analysis of Microarrays | 66.8% (63.9%, 69.6%) | 0.628 |
| XGBoost – TCGA SNP Cohort | 81.1% (78.7%, 83.4%) | 0.787 |
| XGBoost – TCGA WES Cohort | 78.6% (76.0%, 81.0%) | 0.759 |

<sup>a</sup>Models were applied to TCGA SNP cohort unless otherwise specified.
<sup>b</sup>Overall recall is reported as micro-average across all IntClusts.

**Figure 2.**
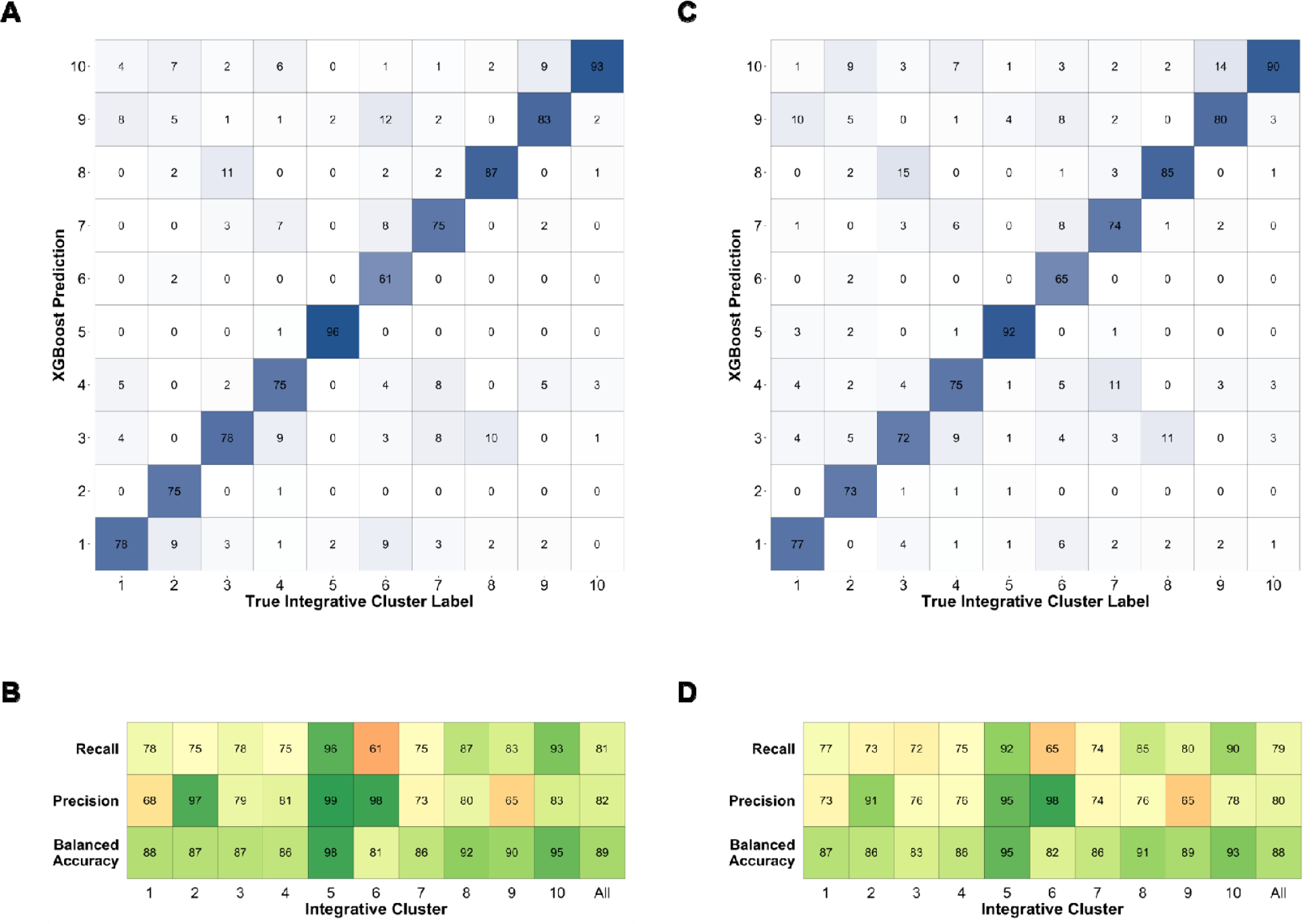
Performance of *CopyClust* on IntClust Label Assignment of TCGA SNP and WES Cohorts. A) Confusion matrix of true IntClust label of TCGA SNP cohort (x-axis) and *CopyClust* prediction (y-axis). Values represent percentage of true IntClust label predicted to be in each class by *CopyClust*. The diagonal represents the percent of samples correctly predicted as a particular IntClust and is equivalent to recall. B) Model performance metrics, where recall = percentage of correctly classified samples per IntClust; precision = percent of correctly classified samples amongst samples predicted as a particular IntClust; and balanced accuracy = mean of recall and specificity. C) Confusion matrix of true IntClust label of TCGA WES cohort (x-axis) and *CopyClust* prediction (y-axis). Values represent percentage of true IntClust label predicted to be in each class by *CopyClust*. The diagonal represents the percent of samples correctly predicted as a particular IntClust and is equivalent to recall. D) Model performance metrics as in B. Overall performance metrics above the “All” column are reported as micro-averages across all IntClusts.

Compared to the current gold standard CN-only *iC10* classifier, *CopyClust* achieved a nine-percentage point greater overall recall when applied to METABRIC (82% vs. 73%) and a 22% and 20% greater overall recall when applied to the TCGA SNP and WES datasets, respectively (81% and 79% vs. 59%) (**Supplemental Table 1**). This increase in the performance of *CopyClust* compared to the *iC10* classifier can likely be attributed to the dominance of GE features in the selected probes of the *iC10* classifier^7^. Features in the *iC10* classifier were taken from the original IntClust manuscript^1^, and only 38 out of 714 (5.3%) of the probes used are gene CN; therefore, the bulk of the features are GE values. The *iC10* classifier is trained using the prediction analysis of microarrays shrunken centroids approach, which was developed for GE analysis^10^. Rather than using a small subset of CN probes, *CopyClust* was trained using features comprising the entire length of the genome. Many IntClusts have key features of their CN profiles that are characteristic for a given IntClust^6^ (e.g. IntClust 5 [chromosome 17q12 amplification] and IntClust 6 [chromosome 8p12 amplification]). The CN probes used by the *iC10* classifier do cover some of these key regions, but they do not cover the characteristics of the entire CN profile, indicating the superiority of *CopyClust* in the absence of GE profiling.

Intricacies of the specific datasets used to train and validate *CopyClust* somewhat limit its generalizability. METABRIC did not set a minimum tumor cellularity^1^, while TCGA set a minimum of 60%^11^. The TCGA cohort may be composed of tumors with a greater average percentage of neoplastic cells; this difference may also account for the stronger signal observed in the TCGA CN profiles relative to the METABRIC CN profiles (**Supplementary Figure S18**), which necessitated feature scaling before model training. Additionally, the need to apply feature scaling across samples limits performance when there are only a single or few samples to classify. Manual curation of genomic ranges developed from PCF was only performed to ensure that ranges did not span multiple chromosomes but ranges still cover regions of telomeres and centromeres. Finally, *CopyClust* was only trained using a single cohort and validated externally on a single cohort, therefore, replication on additional datasets may further improve performance.

*CopyClust* provides an accurate and easily implementable framework for IntClust classification using CN data and achieves a nine-percentage point or greater improvement in overall classification recall compared to the current gold standard approach. Furthermore, *CopyClust* can flexibly handle missing features, is agnostic to differences in genomic profiling platforms, and is easily implementable in an open-source environment, allowing for seamless application to external genomic datasets. The *CopyClust* R package is currently available for download on GitHub (https://github.com/camyoung54/CopyClust).

## Acknowledgements

This research was supported with funding from Cancer Research UK (Caldas Core Grant A16942 and CRUK Cambridge Institute Core Grant A29580), and an ERC Advanced Grant to C.C. from the European Union’s Horizon 2020 research and innovation programme (ERC-2015-AdG-694620). C.C.Y. was supported by the Winston Churchill Foundation of the United States. O.M.R. is supported by NIHR Cambridge Biomedical Research Centre (BRC-1215-20014) and the Medical Research Council (United Kingdom; MC_UU_00002/16). The funder played no role in study design, data collection, analysis and interpretation of data, or the writing of this manuscript. The results shown here are in whole or part based upon data generated by the TCGA Research Network: https://www.cancer.gov/tcga. Figures were created with biorender.com.

