## Supplementary Material for "Development and validation of a reliable DNA copy-number-based machine learning algorithm (*CopyClust*) for breast cancer integrative cluster classification"

**Table of Contents:**

| **Title** | **Pages** |
| --- | --- |
| **Table S1** – Summary of overall model performance metrics | 3 |
| **Table S2** – Genomic regions identified from PCF in hg18, hg19, and hg38 using METABRIC cohort | 4 |
| **Table S3** – Range and optimized values of multiclass XGBoost hyperparameters | 5 |
| **Table S4** – Range of XGBoost hyperparameters investigated for combined integrative clusters 1 and 5 binary models and optimized values | 6 |
| **Table S5** – Range of XGBoost hyperparameters investigated for combined integrative clusters 3 and 8 binary models and optimized values | 6 |
| **Table S6** – Range of XGBoost hyperparameters investigated for combined integrative clusters 4 and 7 binary models and optimized values | 7 |
| **Table S7** – Range of XGBoost hyperparameters investigated for combined integrative clusters 9 and 10 binary models and optimized values | 7 |
| **Figure S1** – Distribution of integrative clusters between METABRIC training and validation cohorts | 8 |
| **Figure S2** – Distribution of integrative clusters between METABRIC and TCGA cohorts | 9 |
| **Figure S3** – PCA of genomic features comparing METABRIC and TCGA | 9 |
| **Figure S4** – Example mean copy number profile prior to fitting PCF to data | 10 |
| **Figure S5** – Mean copy number profile of METABRIC integrative cluster 1 samples with PCF segmentation | 10 |
| **Figure S6** – Mean copy number profile of METABRIC integrative cluster 2 samples with PCF segmentation | 11 |
| **Figure S7** – Mean copy number profile of METABRIC integrative cluster 3 samples with PCF segmentation | 11 |
| **Figure S8** – Mean copy number profile of METABRIC integrative cluster 4 samples with PCF segmentation | 12 |
| **Figure S9** – Mean copy number profile of METABRIC integrative cluster 5 samples with PCF segmentation | 12 |
| **Figure S10** – Mean copy number profile of METABRIC integrative cluster 6 samples with PCF segmentation | 13 |
| **Figure S11** – Mean copy number profile of METABRIC integrative cluster 7 samples with PCF segmentation | 13 |
| **Figure S12** – Mean copy number profile of METABRIC integrative cluster 8 samples with PCF segmentation | 14 |
| **Figure S13** – Mean copy number profile of METABRIC integrative cluster 9 samples with PCF segmentation | 14 |
| **Figure S14** – Mean copy number profile of METABRIC integrative cluster 10 samples with PCF segmentation | 15 |
| **Figure S15** – Histogram of METABRIC integrative cluster 1 LOF values | 15 |
| **Figure S16** – Histogram of XGBoost model accuracy following 1,000 iterations of grid search hyperparameter optimization | 16 |
| **Figure S17** – Scatterplot of XGBoost model accuracy versus number of rounds | 16 |
| **Figure S18** – Mean copy number profile of integrative cluster 1 samples from different cohorts | 17 |
| **Supplemental Methods** | 18–28 |
| **References** | 28–30 |

### Supplementary Tables

**Table S1** – Summary of overall model performance metrics

| **Model** | **Overall Recall** | **Overall Precision** | **Overall Balanced Accuracy** | **Matthews Correlation Coefficient** |
| --- | --- | --- | --- | --- |
| *iC10 Classifier – METABRIC cohort* | 73% | 75% | 85% | 0.703 |
| *iC10 Classifier – TCGA SNP cohort* | 59% | 72% | 77% | 0.563 |
| *XGBoost Internal Validation – 10-class* | 79% | 79% | 88% | 0.757 |
| *XGBoost Internal Validation – 6-class with binary reclassification* | 82% | 82% | 90% | 0.797 |
| *XGBoost External Validation (TCGA SNP) – 10-class* | 76% | 77% | 86% | 0.729 |
| *XGBoost External Validation (TCGA SNP) – 6-class with binary reclassification* | 81% | 82% | 89% | 0.787 |
| *XGBoost External Validation (TCGA WES) – 10-class* | 75% | 77% | 86% | 0.727 |
| *XGBoost External Validation (TCGA WES) – 6-class with binary reclassification* | 79% | 80% | 88% | 0.759 |

Overall performance metrics are reported as micro-averages across all IntClusts.

#### **Table S2** – Genomic Regions Identified from PCF in hg18, hg19, and hg38 Using METABRIC Cohort

*See attached Excel spreadsheet.*

#### **Table S3** – Range and optimized values of multiclass XGBoost hyperparameters

|  |  | **Internal Validation** | | **External Validation – TCGA WES** | | **External Validation – TCGA SNP** | |
| --- | --- | --- | --- | --- | --- | --- | --- |
| *Parameter* | *Range* | *10-Class Model* | *6-Class Model* | *10-Class Model* | *6-Class Model* | *10-Class Model* | *6-Class Model* |
| objective |  | multi:softmax | multi:softmax | multi:softmax | multi:softmax | multi:softmax | multi:softmax |
| eval_metric |  | mlogloss | mlogloss | mlogloss | mlogloss | mlogloss | mlogloss |
| num_class |  | 10 | 6 | 10 | 6 | 10 | 6 |
| max_depth | [6, 10] | 6 | 10 | 8 | 9 | 8 | 9 |
| eta | [0.01, 0.3] | 0.297 | 0.288 | 0.041 | 0.112 | 0.041 | 0.112 |
| gamma | [0.0, 0.2] | 0.109 | 0.079 | 0.032 | 0.033 | 0.032 | 0.033 |
| subsample | [0.5, 1] | 0.759 | 0.681 | 0.517 | 0.768 | 0.517 | 0.768 |
| colsample_bytree | [0.6, 1] | 0.832 | 0.656 | 0.721 | 0.943 | 0.721 | 0.943 |
| min_child_weight | [1, 12] | 5 | 3 | 12 | 12 | 12 | 12 |
| max_delta_step | [1, 10] | 2 | 1 | 8 | 2 | 8 | 2 |
| nrounds | [1, 312] | 47 | 119 | 70 | 95 | 70 | 95 |

#### **Table S4** – Range of XGBoost hyperparameters investigated for combined integrative clusters 1 and 5 binary models and optimized values

| *Parameter* | *Range* | **Internal Validation** | **External Validation – TCGA WES** | **External Validation – TCGA SNP** |
| --- | --- | --- | --- | --- |
| objective |  | binary:logistic | binary:logistic | binary:logistic |
| eval_metric |  | rsme | rsme | rsme |
| max_depth | [6, 10] | 9 | 8 | 8 |
| eta | [0.01, 0.3] | 0.198 | 0.173 | 0.173 |
| gamma | [0.0, 0.2] | 0.123 | 0.067 | 0.067 |
| subsample | [0.5, 1] | 0.545 | 0.648 | 0.648 |
| colsample_bytree | [0.6, 1] | 0.640 | 0.778 | 0.778 |
| min_child_weight | [1, 12] | 4 | 9 | 9 |
| max_delta_step | [1, 10] | 10 | 8 | 8 |
| nrounds | [1, 312] | 28 | 64 | 64 |

**Table S5** – Range of XGBoost hyperparameters investigated for combined integrative clusters 3 and 8 binary models and optimized values

| *Parameter* | *Range* | **Internal Validation** | **External Validation – TCGA WES** | **External Validation – TCGA SNP** |
| --- | --- | --- | --- | --- |
| objective |  | binary:logistic | binary:logistic | binary:logistic |
| eval_metric |  | rsme | rsme | rsme |
| max_depth | [6, 10] | 6 | 8 | 8 |
| eta | [0.01, 0.3] | 0.099 | 0.098 | 0.098 |
| gamma | [0.0, 0.2] | 0.145 | 0.158 | 0.158 |
| subsample | [0.5, 1] | 0.532 | 0.607 | 0.607 |
| colsample_bytree | [0.6, 1] | 0.607 | 0.638 | 0.638 |
| min_child_weight | [1, 12] | 6 | 6 | 6 |
| max_delta_step | [1, 10] | 8 | 4 | 4 |
| nrounds | [1, 312] | 73 | 42 | 42 |

#### **Table S6** – Range of XGBoost hyperparameters investigated for combined integrative clusters 4 and 7 binary models and optimized values

| *Parameter* | *Range* | **Internal Validation** | **External Validation – TCGA WES** | **External Validation – TCGA SNP** |
| --- | --- | --- | --- | --- |
| objective |  | binary:logistic | binary:logistic | binary:logistic |
| eval_metric |  | rsme | rsme | rsme |
| max_depth | [6, 10] | 6 | 7 | 7 |
| eta | [0.01, 0.3] | 0.074 | 0.283 | 0.283 |
| gamma | [0.0, 0.2] | 0.159 | 0.162 | 0.162 |
| subsample | [0.5, 1] | 0.529 | 0.787 | 0.787 |
| colsample_bytree | [0.6, 1] | 0.789 | 0.742 | 0.743 |
| min_child_weight | [1, 12] | 11 | 10 | 10 |
| max_delta_step | [1, 10] | 2 | 6 | 6 |
| nrounds | [1, 312] | 16 | 55 | 55 |

**Table S7** – Range of XGBoost hyperparameters investigated for combined integrative clusters 9 and 10 binary models and optimized values

| *Parameter* | *Range* | **Internal Validation** | **External Validation – TCGA WES** | **External Validation – TCGA SNP** |
| --- | --- | --- | --- | --- |
| objective |  | binary:logistic | binary:logistic | binary:logistic |
| eval_metric |  | rsme | rsme | rsme |
| max_depth | [6, 10] | 10 | 7 | 7 |
| eta | [0.01, 0.3] | 0.071 | 0.226 | 0.226 |
| gamma | [0.0, 0.2] | 0.136 | 0.194 | 0.194 |
| subsample | [0.5, 1] | 0.693 | 0.995 | 0.995 |
| colsample_bytree | [0.6, 1] | 0.681 | 0.800 | 0.800 |
| min_child_weight | [1, 12] | 5 | 11 | 11 |
| max_delta_step | [1, 10] | 4 | 4 | 4 |
| nrounds | [1, 312] | 61 | 64 | 64 |

### Supplementary Figures

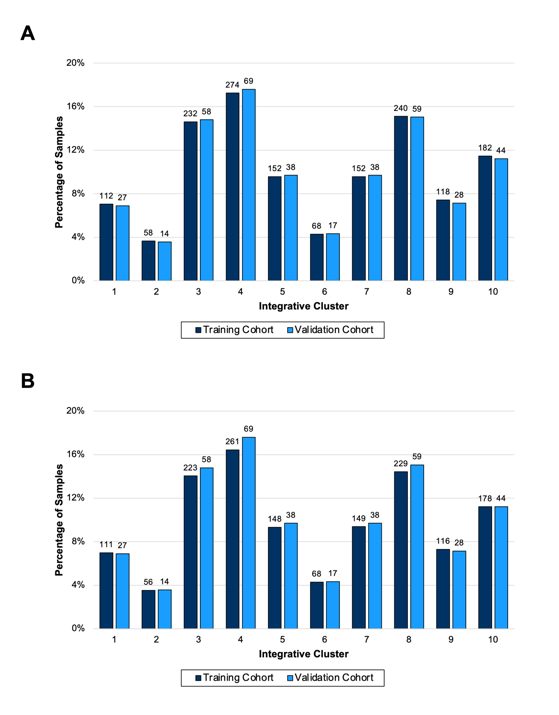

**Figure S1** – Distribution of integrative clusters between METABRIC training and validation cohorts. A) Prior to outlier removal and B) after outlier removal.

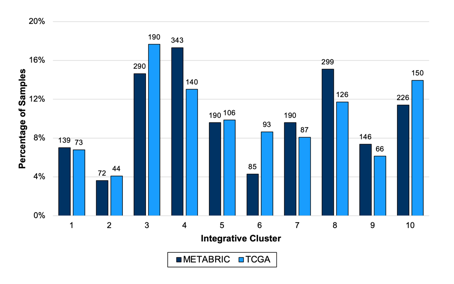

**Figure S2** – Distribution of integrative clusters between METABRIC and TCGA cohorts. Integrative cluster label was assigned from the original manuscript for METABRIC samples and from the results of the combined copy number and gene expression *iC10* classifier for TCGA samples.

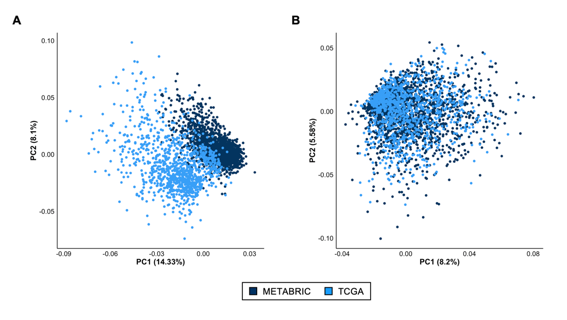

**Figure S3** – PCA of genomic features comparing METABRIC and TCGA. A) Before applying z-score scaling and B) after applying z-score scaling.

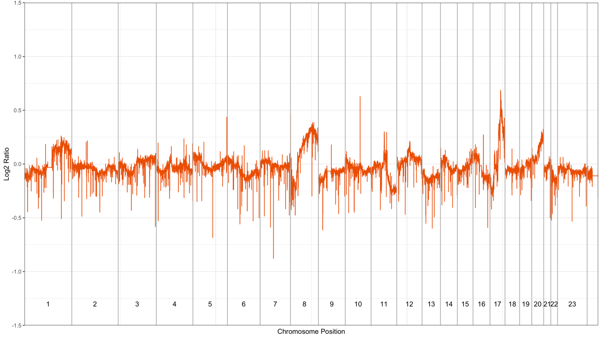

#### **Figure S4** – Example mean copy number profile prior to fitting PCF to data.

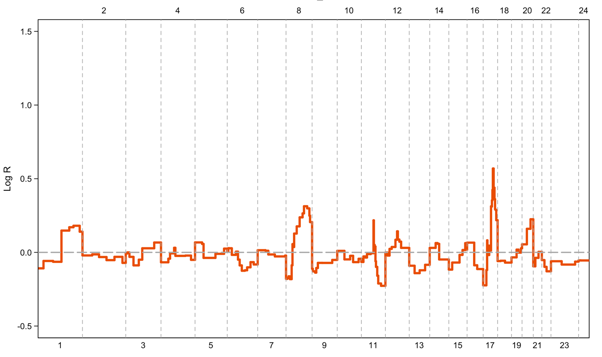

**Figure S5** – Mean copy number profile of METABRIC integrative cluster 1 samples (n=139) with PCF segmentation.

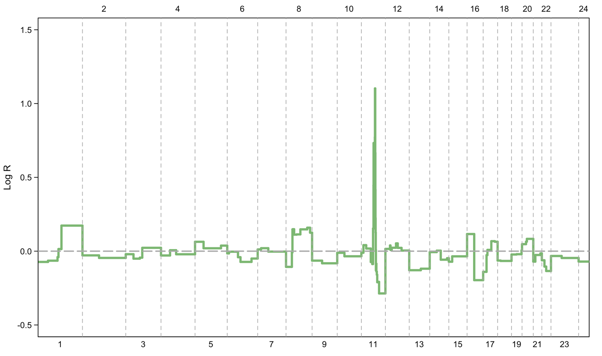

**Figure S6** – Mean copy number profile of METABRIC integrative cluster 2 samples (n=72) with PCF segmentation.

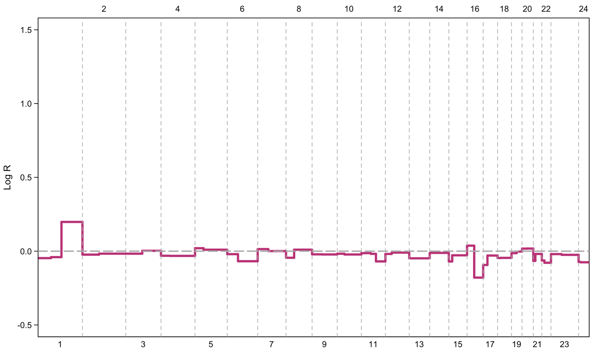

**Figure S7** – Mean copy number profile of METABRIC integrative cluster 3 samples (n=290) with PCF segmentation.

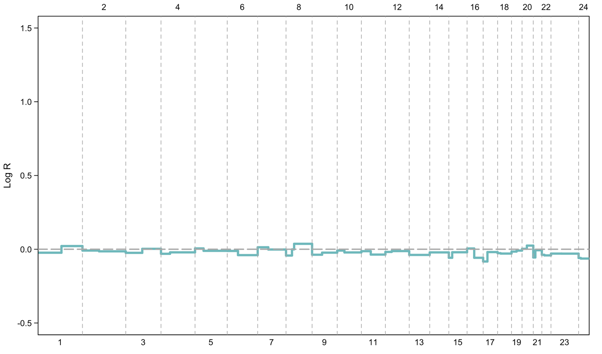

**Figure S8** – Mean copy number profile of METABRIC integrative cluster 4 samples (n=343) with PCF segmentation.

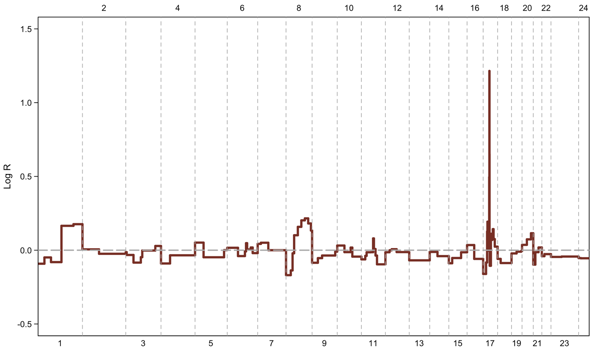

**Figure S9** – Mean copy number profile of METABRIC integrative cluster 5 samples (n=190) with PCF segmentation.

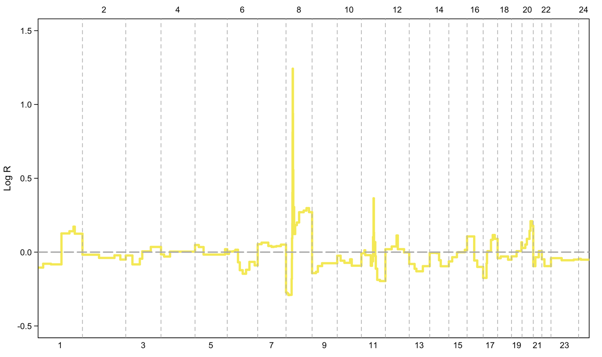

**Figure S10** – Mean copy number profile of METABRIC integrative cluster 6 samples (n=85) with PCF segmentation.

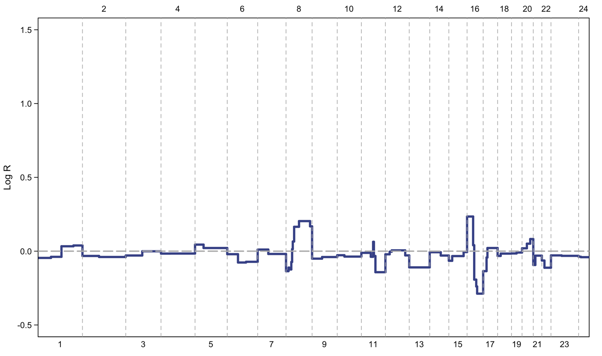

**Figure S11** – Mean copy number profile of METABRIC integrative cluster 7 samples (n=190) with PCF segmentation.

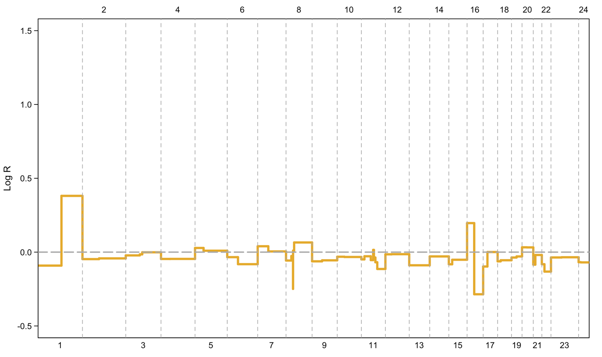

**Figure S12** – Mean copy number profile of METABRIC integrative cluster 8 samples (n=299) with PCF segmentation.

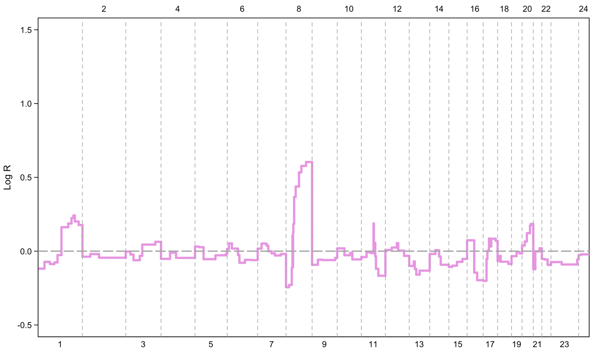

**Figure S13** – Mean copy number profile of METABRIC integrative cluster 9 samples (n=146) with PCF segmentation.

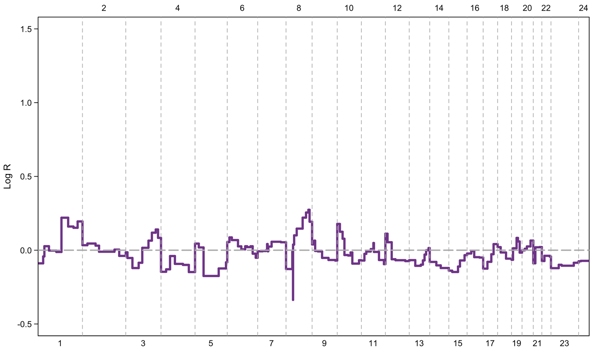

**Figure S14** – Mean copy number profile of METABRIC integrative cluster 10 samples (n=226) with PCF segmentation.

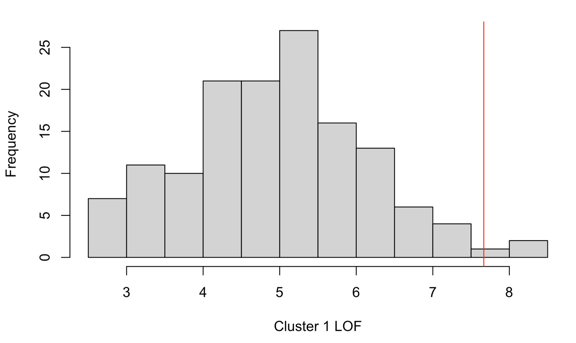

**Figure S15** – Histogram of METABRIC integrative cluster 1 LOF values with the outlier cutoff as vertical red line (cutoff of 7.67).

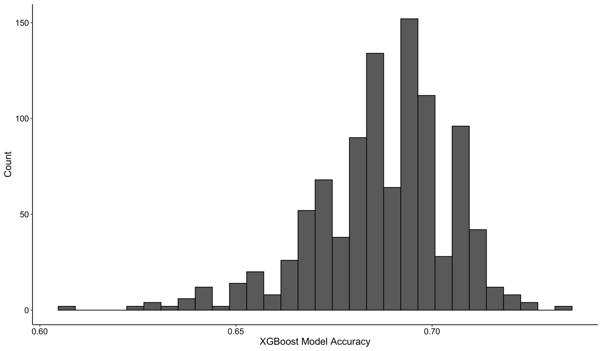

#### **Figure S16** – Histogram of XGBoost model accuracy following 1,000 iterations of random search hyperparameter optimization.

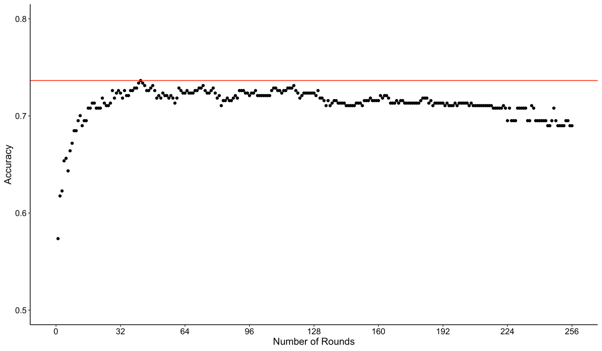

**Figure S17** – Scatterplot of XGBoost model accuracy versus number of rounds (*nrounds* parameter). Red horizontal line represents maximum accuracy value.

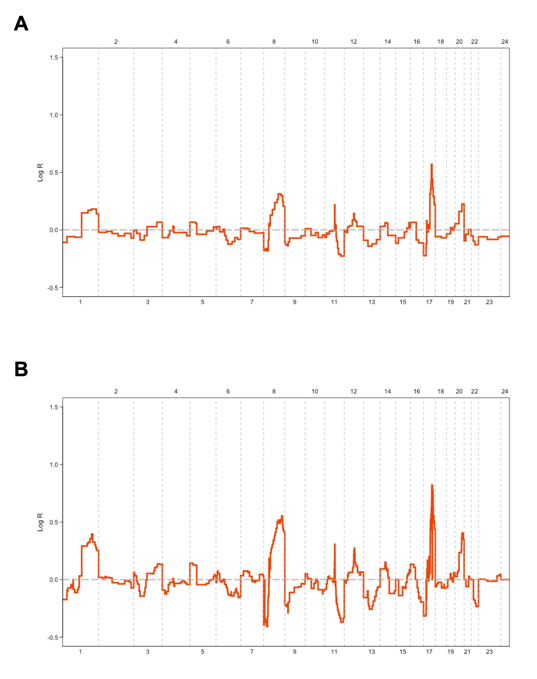

#### **Figure S18** – Mean copy number profile of integrative cluster 1 samples from different cohorts: A) METABRIC and B) TCGA.

**Supplemental Methods:**

### *Training and Validation Cohorts*

This analysis used the previously published data from METABRIC cohort consisting of over 2,000 primary breast cancer samples from tumor banks in the United Kingdom and Canada^1^ as the primary dataset. As previously described DNA and RNA were isolated from the tumor samples and hybridized to the Affymetrix SNP Array 6.0 and Illumina HT-12 v3 platforms for genomic and gene expression profiling, respectively. The original IntClusts were developed using a subset of these samples and consisted of a training cohort of 997 samples and a validation cohort of 995 samples^1^. For this analysis, the original METABRIC training and validation cohorts were grouped into a single cohort and 12 replicate samples were removed, resulting in an analysis cohort of 1,980 samples. DNA copy number, RNA gene expression, and relevant clinical data including IntClust label, estrogen receptor (ER) status, and tumor purity are publicly available online and were downloaded from cBioPortal^2-4^.

The TCGA dataset^5,6^, a publicly available dataset developed by the National Cancer Institute (NCI) in the United States, was used for external validation in this analysis. Among all samples from this cohort, 1,075 primary breast cancer samples with available DNA copy number and RNA gene expression data were selected for analysis. RNA gene expression profiling was performed on the Illumina HiSeq 2000 platform and downloaded from cBioPortal^7^. Two genomic profiling methods were investigated: Affymetrix SNP 6.0 data were downloaded from the NCI Genomic Data Commons^8,9^ and whole-exome sequencing (WES) data from Illumina HiSeq 2000 platform were downloaded from cBioPortal^7^.

### *Dataset Format*

The Affymetrix SNP Array 6.0 platform used for the TCGA cohort contained 1,876,300 probes, with half distributed across the genome for CN profiling and the other half at locations of known SNPs. Pre-processed copy number data were downloaded and expressed as log2 segmented means in the format produced by the *DNAcopy* R package (v1.7)^10^. The Illumina HT-12 v3 platform used for METABRIC consisted of 48,804 probes across more than 25,000 genes. The Illumina HiSeq 2000 platform used for TCGA RNA and WES generated two billion paired end reads and 200 gigabases of filtered data. METABRIC^1^ and TCGA^11^ data were preprocessed according to individual study protocols.

### *Partition of METABRIC Dataset into Training and Validation Cohorts*

The 1,980 METABRIC samples were divided into two groups: a training cohort consisting of 80% of samples (n=1,588) and a validation cohort of the remaining 20% (n=392) using the *createDataPartition* function with default parameters from the *caret* R package (v6.0-93)^12^ with an equal distribution of IntClusts in each cohort (**Supplementary Figure S1**). The validation samples were completely held-out from the full training set and used to assess model performance for internal validation. Subsequently, these samples were included in the full training set to develop the models for external validation and performance was assessed using TCGA datasets.

### *Integrative Cluster Label Assignment*

For samples from the METABRIC cohort, the IntClust label was assigned from the original IntClust classifier as reported in cBioPortal^1,2,13^ and used as the gold standard label for analysis. For samples from the TCGA dataset, the IntClust label was assigned using the combined copy number and gene expression classifier from the *iC10* R package (v1.5)^14^ using data generated from the arrays. Copy number and gene expression features were matched using the *matchFeatures* function, and gene expression data was then normalized using the *normalizeFeatures* function using default parameters. The output from *matchFeatures* and *normalizeFeatures* was used to determine the IntClust label using the *iC10* function using default parameters. The gold standard IntClust label for TCGA samples was assigned using the output from this combined DNA copy number and RNA gene expression *iC10* classifier.

### *Development of Feature Set using Piecewise Constant Fits*

Genomic features for analysis were selected by first separating training samples into 10 groups based on IntClust label. **Supplementary Figures S5–S14** highlight the differences in copy number landscapes amongst the subtypes. In these figures, the mean copy number log2 ratio was calculated at the 1,876,300 probe positions among the samples within each IntClust group and plotted against genomic position to allow for the visualization of the average copy number profile for each IntClust. An example copy number profile is found in **Supplementary Figure S4**. To reduce noise amongst the copy number profiles and simplify analysis by reducing the number of features, a piecewise constant fit (PCF) was applied to the copy number profiles of all samples within an IntClust group using the *copynumber* R package (v1.38.0)^15,16^. The *copynumber* package fits a PCF to the data through penalized least squares minimization^15^ to locate areas or segments of equal copy numbers by minimizing the distance between the PCF and the observed data, while imposing a penalty for each discontinuity in the PCF^16^. The *multipcf* function was applied to all samples in each IntClust with “normalize” set to FALSE and otherwise default parameters. This process generated 10 individual PCFs, one per IntClust, each with unique genomic breakpoints informed by the characteristics of each IntClust copy number profile.

Unique breakpoints across all IntClust copy number profiles were identified using the *GenomicRanges* R package (v1.50.2)^17^. Iterating from chromosome 1, each time a breakpoint was identified in one of the IntClust PCFs, a new genomic range was defined. This method allowed for the preservation of the unique characteristics of each individual IntClust copy number profile, which differed throughout the genome. This process was continued for the entirety of the genome and 478 genomic ranges were identified. Genomic ranges did not span multiple chromosomes. Informed by the genomic start and end positions of each range, probe positions were mapped to an individual range. Then, the mean log2 copy number value was calculated from the probes within a given range for each sample in both the training and validation cohorts. These mean log2 copy number values were the features used for training and validation of the machine learning models.

Feature generation was completed independently using the 1,588 samples in the METABRIC training cohort for internal validation and later using all 1,980 METABRIC samples for external validation. The genomic ranges determined from the 1,588 sample METABRIC training cohort were used to calculate the mean log2 copy number values for the METABRIC training and validation cohorts as features for internal validation. Separately, the genomic ranges determined from the entire 1,980 sample METABRIC cohort were used to calculate the mean log2 copy number value for all METABRIC samples and for both the TCGA SNP and WES datasets as features for external validation. Plots of the average copy number profile for each IntClust using all 1,980 samples from the METABRIC cohort can be found in the **Supplementary Figures S5–S14**.

### *Conversion of hg18 Probe Positions to hg19 and hg38*

Genomic ranges were generated from the probe positions of the Affymetrix SNP Array 6.0, which were originally mapped to the hg18 reference genome for the METABRIC cohort. The TCGA WES dataset was mapped to hg19, while the TCGA SNP dataset was mapped to hg38. To convert probe positions from hg18 to hg19 and from hg18 to hg38, each hg18 probe position from the METABRIC Affymetrix SNP Array 6.0 was uploaded to *liftOver*^18^ and mapped to either the hg19 or hg38 reference genomes. Using the output from *liftOver*, the same number and sequential position of probes were assigned to genomic ranges, using the hg19 and hg38 reference genome locations as the start and end locations of the new genomic ranges (**Supplementary Table S2**). Genomic ranges that spanned multiple chromosomes following *liftOver* were amended so that the upper bound of the range corresponded to the last base pair of the starting chromosome for a given reference genome.

### *Evaluation and Removal of Outliers Using Local Outlier Factors*

Before the development of machine learning models, sample outliers from each IntClust group were identified and removed from the training dataset. Principal component analysis (PCA) was performed on the features of all samples from a given IntClust within the training dataset. PCA was performed using the *prcomp* function with default parameters. A local outlier factor (LOF)^19^ was computed for each sample based on its position within the PCA coordinate space. The LOF was calculated by summing the Euclidian distance between the sample and its *k* nearest neighbors and dividing by *k*. This process was repeated for each subspace within the PCA. The value of *k* was defined as 20% of the number of samples within an IntClust group, rounded to the nearest whole number. The *k* nearest neighbors were identified using the *get.knn* function with default parameters from the *FNN* R package (v1.1.3.1)^20^. LOFs for each PCA subspace were weighted by multiplying the value by the sum of the variance explained by the components bounding the coordinate space. A single LOF was then calculated for each sample by summing the LOF values from all PCA subspaces. This calculation is defined mathematically by the formula given in Equation 1.

${LOF}_{i}=\sum_{j=1}^{F} w_{j}\sum_{k\in nn({PC}_{j})} \frac{d({PC}_{i,j},{PC}_{k,j})}{\#k}$	(1)

Where:

*i* is a sample

*j* is a dimension of the PC

*k* is a nearest neighbor of sample *i*

F is the number of PC dimensions

PC_ij_ is the projection of sample *i* in PC *j*

W_j_ is the weight (variance of PC_j_)

The LOFs from each IntClust group were visualized as a histogram and assessed for normality using a Shapiro-Wilk test using the *shapiro.test* function with default parameters. Since the distributions were considered to be normal, outliers were defined as being greater than 1.5 times the interquartile range (IQR) plus the 75^th^ percentile of the distribution^21^, as defined by Equation 2.

$LOF\geq1.5*IQR+75th percentile$	(2)

Samples meeting this definition were removed from the training dataset before model development. This process was repeated individually using the 1,588 METABRIC samples in the training cohort for internal validation and using all 1,980 METABRIC samples for external validation. An example histogram of the distribution of LOFs of METABRIC IntClust 1 samples and the outlier cutoff value can be found in the **Supplementary Figure S15**.

### *Training and Optimization of XGBoost Machine Learning Models*

XGBoost modeling was performed using the framework provided by the *xgboost* R package (v1.7.3.1)^22^. Training data were formatted such that rows corresponded to individual samples while columns corresponded to genomic range features tabulated with average log2 copy number values. Two model approaches were implemented: multiclass (a prediction with more than two possible classes) and binary (a prediction with exactly two possible classes).

To perform hyperparameter optimization, the XGBoost machine learning models were subjected to stratified fivefold cross-validation in which 20% of the training dataset was excluded from each fold and used as validation. Each sample only appeared in a single fold and each fold contained an equal distribution of IntClusts. These folds were iterated through treating each one as the validation set in each iteration, with the remaining four folds combined as the training set. The performance of the hyperparameters was assessed using the independent, unseen, held-out validation fold. These iterations were repeated for various sets of hyperparameters selected via random search optimization, which performs better than grid search or manual search optimization^23^. All iterations were performed using a constant, randomized seed as set by the *set.seed* function. The hyperparameters that resulted in the lowest mean objective value (log-loss score for multiclass models and root square mean error for binary models) were selected for use in the final models and these models were trained using the entire training dataset. Cross-validation was performed using the *xgb.cv* function with “early_stopping_rounds” set to 8 and otherwise default parameters and the model training was completed using the *xgb.train* function with optimized hyperparameters and otherwise default parameters.

Model objectives were set to “multi:softmax” for multiclass models and “binary:logistic” for binary models with “mlogloss” and “rsme” as evaluation metrics, respectively. The IntClust label with the highest probability was assigned automatically using the “multi:softmax” objective in the multiclass models, whereas in the binary models, the lower IntClust label – for example, IntClust 3 when combining IntClust 3 and IntClust 8 – was assigned a value of 0 and the higher IntClust label assigned a value of 1 with the IntClust label assigned according to whether the predicted model probability was closer to 0 or 1. The “multi:softprob” objective was also used for multiclass models to obtain a model probability for all IntClust labels. The model prediction was determined using the *predict* function with default parameters.

Hyperparameters optimized during cross-validation included: “max_depth”, “eta”, “gamma”, “subsample”, “colsample_bytree”, “min_child_weight”, “max_delta_step”, and “nrounds”. All other XGBoost parameters were set to their default. A table of the upper and lower bounds of these hyperparameters used during random search optimization and their optimal values for both multiclass and binary models can be found in the **Supplementary Tables S3–S7**. Hyperparameter ranges were selected on the basis of prior literature^24^. For each model, random search hyperparameter optimization was performed for 1,000 iterations, with the hyperparameters resulting in highest accuracy selected for implementation in the final models. Model accuracy did not vary significantly among different sets of hyperparameters, and this number of iterations appeared to capture the best performing set of hyperparameters. A histogram of model accuracy after 1,000 iterations of random hyperparameters can be found in the **Supplementary Figure S16**. The “nrounds” parameter was optimized individually for each model once the set of hyperparameters was selected. Model accuracy was assessed for all values of “nrounds” between 1 and 256, with the value resulting in the greatest model accuracy selected for implementation in final models. If two values of “nrounds” resulted in the same model accuracy, the lower value was selected to decrease runtime and because increasing the value of “nrounds” tended to result in overfitting and a decrease in model accuracy (**Supplementary Figure S17**). As mentioned, early stopping was also applied to reduce overfitting.

### *Internal and External Validation of XGBoost Machine Learning Models*

A 10-class multiclass model to predict all 10 IntClust groups individually was trained initially. As a reduction in the number of classes and combination of similar classes in a multiclass model has been shown to increase performance^25^, and multiple binary models tend to perform better than multiclass models^26^, four pairs of IntClusts with similar mean copy number profiles were combined and a 6-class multiclass model was trained. These pairs consisted of: IntClusts 1 and 5, IntClusts 3 and 8, IntClusts 4 and 7, and IntClusts 9 and 10. Rather than predicting each IntClust individually, the 6-class model predicted that a sample belonged to one of the following groups: IntClust 1 or 5, IntClust 2, IntClust 3 or 8, IntClust 4 or 7, IntClust 6, or IntClust 9 or 10. To reclassify the pairs of IntClusts into all 10 IntClusts, binary models were trained and optimized using only samples from the training cohort that belonged to the two IntClust groups of interest. Following the training and optimization process, four binary reclassification models were generated using the training cohort. Then, samples predicted to be in one of the 4 combined IntClust groups from the 6-class model were reassigned to a single IntClust label using the corresponding binary classifier, resulting in a discrete IntClust classification for all samples. Model performance was ascertained based on the accuracy of prediction for both the 10-class model and the 6-class model with binary model reclassification.

Internal validation was performed solely using the METABRIC cohort data. The internal validation models were trained and optimized using the METABRIC training cohort described above**,** and validated using the corresponding validation cohort. To validate the performance of the XGBoost model approach on both a different cohort and a different genomic profiling platform, external validation was performed using the TCGA SNP (cohort difference only) and TCGA WES (cohort and platform difference) datasets. In both external validation cases, the models were trained and optimized using the entire METABRIC dataset and validated on the entire TCGA SNP and TCGA WES datasets, respectively.

### *Scaling of Features for External Datasets*

To adjust for differences in the signal to noise of log2 ratios between samples from the METABRIC and TCGA datasets, feature values were scaled prior to model training and optimization during external validation. Z-score scaling was performed using the *scale* function for each individual feature with mean set to 0 and standard deviation set to 1. Scaling of features significantly reduced intra-cohort feature differences as visualized by PCA (**Supplementary Figure S3**).

### *Model Performance Benchmarking*

To assess the performance of the XGBoost models relative to other machine learning algorithms, four alternative multiclass classifiers were trained and optimized using the scaled features from the entire METABRIC cohort and their performance was assessed using the scaled features from the entire TCGA SNP cohort (**Supplementary Table S8**). Model hyperparameter optimization was performed using the same methodology above. Overall multiclass classification accuracy was the metric used to assess performance. The four model techniques used as comparison were random forest, support vector machine (SVM), Light Gradient Boosting Machine (LightGBM)^27^, and PAM. The random forest model was implemented using *randomForest* function from the *randomForest* R package (v4.7.1.1)^28^ with the optimized parameters “ntrees” and “mtry”. SVM was implemented using the *e1701* R package (v1.7.12)^29^. The model was built with “type” = “C-classification” and “kernel” = “radial” as well as optimized hyperparameters. The LightGBM model was built using the framework provided by the *lightgbm* R package (v3.3.5)^30^. Model training was performed using the *lightgbm* function with optimized hyperparameters, “objective” = “multiclass”, and “num_class” = 10. The PAM methodology was implemented using the *pamr* package (v1.56.1)^31^ with the classifier trained using the *pamr.train* function with an optimized “n.threshold” parameter with otherwise default parameters. Overall classifier performance metrics including recall, precision, balanced accuracy, and 95% confidence intervals were determined using the *confusionMatrix* function from the *caret* R package (v6.0-93)^12^ with default parameters. Since it is considered to be a more robust model performance metric than accuracy, precision, and balanced accuracy, particularly among unbalanced datasets, the Matthews Correlation Coefficient^32^ was also calculated using the *mcc* function from the *mltools* R package (v0.3.5)^33^.

### *Definition of Performance Metrics*

Model performance was assessed for both the 10-class model and the 6-class model with binary model reclassification for internal validation and both cases of external validation using standard multiclass model performance metrics^34^. Micro-weighted averages, the summation of values across all classes, were calculated for all summary performance metrics. Performance was also assessed by IntClust. Overall recall was the primary performance metric of interest. Balanced accuracy is also reported to account for the difference in the population prevalence of IntClusts. Performance metrics were defined as follows:

- Overall recall is defined as the sum of true positives (TP) and true negatives (TN) divided by the sum of TP, TN, false positives (FP), and false negatives (FN).

$Overall Recall=\frac{TP+TN}{TP+TN+FP+FN}$	(3)

- Recall is defined as the TP divided by the sum of TP and FN

$Recall=\frac{TP}{TP+FN}$	(4)

- Precision is defined as the TP divided by the sum of TP and FP

$Precision=\frac{TP}{TP+FP}$	(5)

- Balanced accuracy is defined as the arithmetic mean of sensitivity (referred to herein as recall) and specificity (TN divided by the sum of TN and FP)

$Balanced Accuracy=\frac{\frac{TP}{TP+FN}+\frac{TN}{TN+FP}}{2}$	(6)

- Matthews Correlation Coefficient (MCC) is defined as:

$MCC=\frac{TN\times TN-FP\times FN}{\sqrt{(TP+FP)(TP+FN)(TN+FP)(TN+FN)}}$	(7)
